# DYN-1 regulates SPD-2 and PLK-1 localization and mitotic spindle pole organization

**DOI:** 10.1101/2025.07.21.665896

**Authors:** Carter Dierlam, Stephanie Held, Jiya Hastings, Livinus Anyanwu, Robert H. Newman, Jyoti Iyer

**Affiliations:** Department of Cell Biology and Molecular Genetics, University of Maryland College Park, College Park, MD 20742; Department of Biology, North Carolina Agricultural and Technical State University, Greensboro, NC 27411; Department of Chemistry and Biochemistry, University of Tulsa, Tulsa, OK 74104

**Author notes:** To whom correspondence should be addressed: Jyoti Iyer.

**Keywords:** dynamin, PLK-1, SPD-2, mitosis, midbody, centrosome

## Abstract

Accurate mitosis is essential for genome stability. Although dynamin is best known for its role in vesicular trafficking, its functions during mitosis remain unclear. Here, we identify a mitotic role for the *C. elegans* dynamin homolog DYN-1 in regulating spindle assembly and the localization of the key mitotic regulators SPD-2 and PLK-1. Depletion of DYN-1 results in enlarged metaphase spindle poles and increased centrosomal SPD-2 and PLK-1 levels. These effects likely depend on DYN-1-mediated vesicular trafficking, as shown by assays with the *dyn-1(ky51)* allele. Additionally, a subset of DYN-1-depleted embryos exhibit altered PLK-1 localization at the midbody during late mitosis, correlating with midbody assembly defects. Together, these findings establish DYN-1 as a previously unrecognized regulator of spindle pole organization and early midbody assembly.

**SIGNIFICANCE STATEMENT:** Dynamin is primarily known for its function in endocytosis, but its involvement in mitosis is not well defined. This study shows that dynamin helps organize mitotic spindle poles, ensures proper centrosome localization of SPD-2 and PLK-1, and supports early steps of midbody assembly in *C. elegans* embryos. These findings reveal a previously unrecognized mitotic role for dynamin and suggest new links between membrane dynamics, spindle organization, and the cytokinetic machinery.

## INTRODUCTION

Accurate cell division is essential for proper chromosome segregation, and mitotic errors can lead to aneuploidy—a distinguishing feature of many cancers (Levine and Holland, 2018). Understanding the molecular mechanisms that regulate mitosis is therefore critical. The GTPase dynamin is a large ∼100 kDa protein that is best known for its role in the scission of endocytic vesicles (Hinshaw, 2000; Praefcke and McMahon, 2004). *C. elegans* possesses a single classical dynamin encoded by the *dyn-1* gene (Clark *et al*., 1997). Interestingly, some of the earliest studies identified dynamin as a microtubule-binding protein (Paschal *et al*., 1987; Shpetner and Vallee, 1989). However, dynamin’s function in microtubule-associated processes such as spindle assembly and midbody organization during mitosis is not well understood. While DYN-1 was shown to localize to the midbody (Thompson *et al*., 2002), and mediate membrane remodeling during the late stages of abscission (König *et al*., 2017), its potential role in early midbody assembly remains unexplored.

The coiled-coil protein SPindle Defective 2 (SPD-2) (the *C. elegans* homolog of human CEP192) is essential for proper centrosome biogenesis and spindle assembly during mitosis (O’Connell *et al*., 2000; Kemp *et al*., 2004; Pelletier *et al*., 2004; Gomez-Ferreria *et al*., 2007, Zhu *et al*., 2008). SPD-2 levels at the centrosomes are regulated by proteins including SAS-7, SPD-5, DHC-1 and AIR-1, among others (Sugioka *et al*., 2017; Kemp *et al*., 2004; Pelletier *et al*., 2004). Proper levels of centrosome-associated SPD-2 are required for proper centriole duplication, pericentriolar material (PCM) organization and spindle assembly (O’Connell *et al*., 2000; Kemp *et al*., 2004; Pelletier *et al*., 2004). SPD-2 directly binds to the kinase Polo-like kinase 1 (PLK-1) to recruit the kinase to the centrosomes (Boxem *et al.,* 2008; Decker *et al.,* 2011).

PLK1 is a mitotic master regulator that is essential for several mitotic processes including mitotic entry, centrosome dynamics, chromosome congression, chromosome alignment and cytokinesis (Petronczki *et al.,* 2008; Archambault and Glover, 2009; Zitouni *et al.,* 2014). PLK1 exhibits a dynamic localization pattern at different stages of mitosis. During early mitosis, PLK1 localizes prominently to the centrosomes, and the kinetochores. At anaphase, it re-localizes to the spindle midzone and at telophase, it becomes enriched at the midbody (Golsteyn *et al.,* 1995; Lee *et al.,* 1995; Arnaud *et al.,* 1998; Barr *et al*., 2004; Petronczki *et al.,* 2008; Schmucker and Sumara, 2014). The timely localization of PLK1 to various cellular components is essential for its different important roles during mitosis. Importantly, removal of PLK1 from the centrosomes is required for proper mitotic progression (Kishi *et al.,* 2009). Although the kinase activity of PLK1 is thought to be required for PLK1 removal from the centrosomes (Kishi *et al.,* 2009), the mechanism by which PLK1 is removed from the centrosomes remains unclear.

## RESULTS AND DISCUSSION

### *dyn-1* depletion results in abnormal metaphase spindle pole assembly and increased levels of SPD-2

To determine the effect of DYN-1 depletion in *C. elegans* embryos, we knocked down endogenous DYN-1 using *RNAi* (Supplemental Figure S1, A and B). Interestingly, DYN-1–depleted embryos (n=20) were noticeably smaller than controls (n=23), exhibiting a reduced average cytoplasmic area (∼906 µm² vs. ∼1200 µm²; p < 0.0001) (Supplemental Figure S1, C and D). DYN-1 was previously shown to localize to the mitotic spindle at metaphase, suggesting a potential role in metaphase spindle assembly (Thompson *et al.,* 2002). To determine the effect of DYN-1 depletion on metaphase spindle assembly, we examined the effect of DYN-1 depletion on spindle pole size using mCherry::tubulin as a proxy (Figure 1). Upon measuring spindle pole area, we determined that DYN-1-depleted embryos (n=58 spindle poles) exhibited larger spindle poles than control (*smd-1(RNAi)*)*-*depleted embryos (n=53 spindle poles; p=0.0018) (Figure 1, A and B). Specifically, the mean spindle pole area of control RNAi embryos was 14.6 µm^2^ while that of *dyn-1(RNAi)* embryos was 17.36 µm^2^. Thus, our data have identified a novel role for the endocytic protein DYN-1 in regulating metaphase spindle pole assembly. An important regulator of spindle pole and centrosome size is the PCM protein SPD-2 (O’Connell *et al*., 2000; Kemp *et al*., 2004; Pelletier *et al*.; Decker *et al.,* 2011). Therefore, we asked whether SPD-2 levels at the centrosomes were altered between control RNAi and *dyn-1(RNAi)* treated embryos. Our analysis indicated that centrosome-associated SPD-2 levels were significantly increased ∼1.7-fold upon DYN-1 depletion (Figure 1, C and D; n=66 for *dyn-1(RNAi)*; n=60 for controls; p < 0.0001).

**FIGURE 1.**
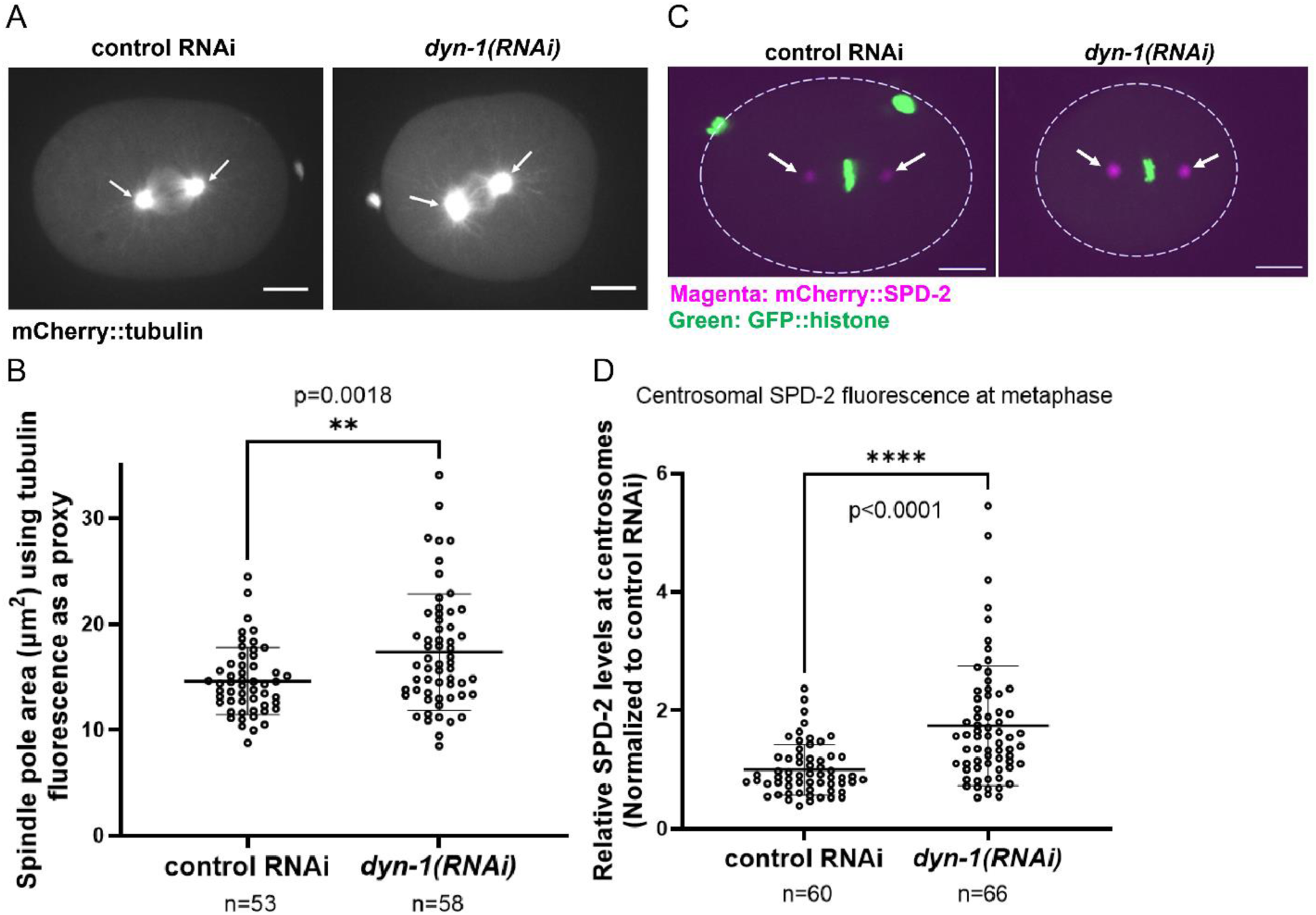
DYN-1 depletion increases spindle pole size and elevates SPD-2 at centrosomes. (A) Spindle pole morphology in embryos expressing mCherry::tubulin, showing enlarged spindle poles upon *dyn-1(RNAi)*. White arrows: spindle poles. (B) Quantification of spindle pole area using mCherry::tubulin fluorescence. Each point = single pole; n = spindle poles analyzed. Error bars = s.d.; middle bar = mean. (C) SPD-2localization at metaphase centrosomes in control and *dyn-1(RNAi)* embryos. Dotted outline demarcates the embryo boundary; white arrows: centrosome-localized SPD-2 (D) Relative centrosomal SPD-2 intensity normalized to control RNAi. n = centrosomes. Error bars = s.d.; middle bar = mean. (B, D): unpaired two-tailed t-test; Experimental replicates: 4 for all assays. All fluorescence images include a 10 µm scale bar.

### DYN-1 depletion increases levels of centrosome-associated PLK-1

Because SPD-2 directly binds to PLK-1 and recruits it to centrosomes (Boxem *et al.,* 2008; Decker *et al.,* 2011), and SPD-2 levels increase upon DYN-1 depletion (Figure 1, C and D), we next asked whether PLK-1 also becomes elevated at centrosomes under these conditions. To determine if DYN-1 depletion affects centrosomal PLK-1, we performed live imaging on a strain co-expressing mCherry::tubulin and CRISPR-tagged endogenous PLK-1::sfGFP (Martino *et al.,* 2017). We then knocked down endogenous DYN-1 in this strain using *RNAi* and performed live imaging of control (*smd-1*(*RNAi)*) and *dyn-1(RNAi)* treated embryos using spinning-disk confocal microscopy. Consistent with our hypothesis, our analysis revealed that PLK-1 levels at the centrosomes were significantly elevated upon DYN-1 depletion (Figure 2, A and B). Specifically, we found that PLK-1 intensity was ∼2.5-fold higher in *dyn-1(RNAi)* embryos (n=52) than controls (n=58; p < 0.0001) (Figure 2, A and B).

**FIGURE 2.**
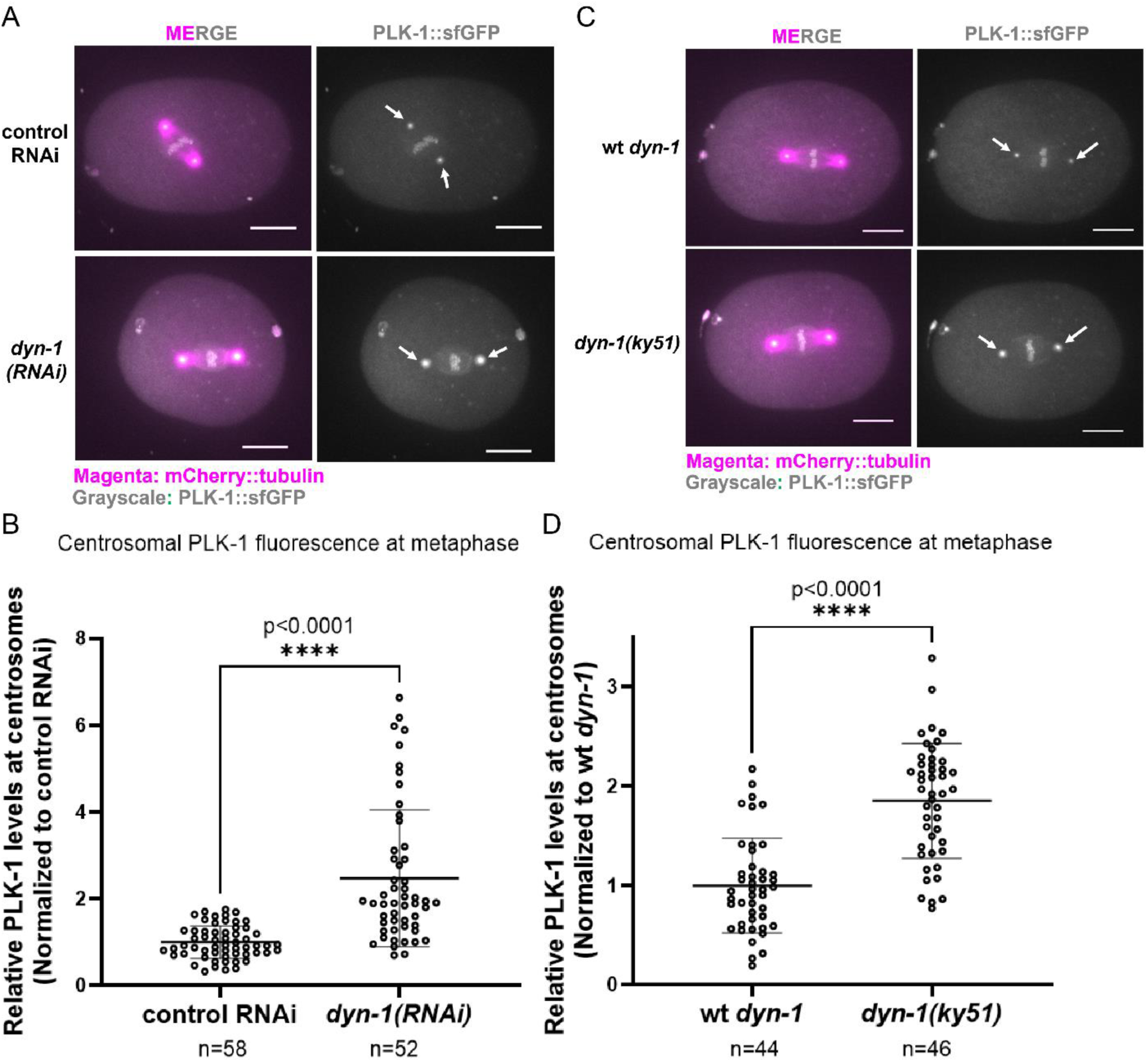
DYN-1 depletion increases centrosomal PLK-1 at metaphase. (A) PLK-1::sfGFP localization at metaphase centrosomes in control RNAi vs *dyn-1(RNAi)* embryos. White arrows: centrosome-localized PLK-1::sfGFP. (B) Quantification of PLK-1::sfGFP centrosomal fluorescence at metaphase. n = centrosomes. Error bars = s.d.; middle bar = mean. (C) PLK-1::sfGFP localization in wild-type (wt) *dyn-1* and *dyn-1(ky51)*. White arrows: centrosome-localized PLK-1::sfGFP. (D) Relative centrosomal PLK-1intensity in wild-type (wt) *dyn-1* and *dyn-1(ky51)*. n = centrosomes. Error bars = s.d.; middle bar = mean. (B, D): unpaired two-tailed t-test; Experimental replicates: 3 for all assays. All fluorescence images include a 10 µm scale bar.

The *dyn-1(ky51)* GTPase mutant, which perturbs vesicular trafficking (Clark *et al*., 1997; Grant and Hirsch, 1999), also exhibited increased centrosomal PLK-1 (Figure 2). Specifically, PLK-1 levels were on average elevated ∼1.85-fold at *dyn-1(ky51)* centrosomes (n=46 centrosomes) as compared with control centrosomes (n=44 centrosomes; p<0.0001) (Figure 2, C and D). These data indicate that the vesicular trafficking function of DYN-1 is required for its regulation of centrosome-associated PLK-1 levels. Furthermore, these data also confirm the specificity of our *dyn-1(RNAi)* phenotype.

Endogenous PLK-1 is a low-abundance protein in wild-type (N2) whole-worm lysates (Chen *et al*., 2023), and the preparation of embryonic extracts is technically labor-intensive. Therefore, to sensitively assess whether total PLK-1 levels differ between control and *dyn-1(ky51)* embryos, we measured PLK-1::sfGFP fluorescence intensity in the anterior and posterior cytoplasm of metaphase embryos (Supplemental Figure S2). We reasoned that if total PLK-1 levels are increased upon impairing DYN-1 function, we should see a proportional increase in cytoplasmic PLK-1 fluorescence in the *dyn-1(ky51)* embryos. Our analysis revealed a modest ∼7% increase in PLK-1 levels in the anterior and posterior cytoplasm of *dyn-1(ky51)* embryos as compared with controls (p=0.0234 (anterior), p=0.0318 (posterior)) (Supplemental Figure S2). Given that PLK-1 levels at metaphase centrosomes increase by ∼1.85-fold in *dyn-1(ky51)* embryos (Figure 2, C and D), this modest ∼1.07-fold increase in cytoplasmic PLK-1 is unlikely to fully explain the elevated centrosome-associated PLK-1 levels observed in these mutants.

### PLK-1 persists at the centrosomes following DYN-1 depletion in a subset of embryos during the later stages of mitosis

PLK-1 exhibits a spatially and temporally regulated localization pattern during different stages of mitosis (Chase *et al.,* 2000; Budirahardja and Gönczy, 2008; Martino *et al.,* 2017). It transitions from its location at the centrosomes and the kinetochores during early mitosis, to the spindle midzone and the midbody during late mitosis (Chase *et al.,* 2000; Budirahardja and Gönczy, 2008; Martino *et al.,* 2017). While performing live imaging, we noticed several embryos where PLK-1 appeared to persist at the centrosomes (specifically at the PCM) during the later stages of mitosis (Figure 3A). To quantify this phenotype, we analyzed PLK-1::sfGFP fluorescence intensity at anterior centrosomes at different cell cycle stages in both control RNAi and *dyn-1(RNAi)* treated embryos (Figure 3; Supplemental Figure S3). Note that we measured PLK-1::sfGFP at anterior centrosomes only because cytoplasmic background is more uniform there; in contrast, the posterior cytoplasm contains punctate, uneven PLK-1 that makes measurements unreliable. Our quantitative analysis revealed that there was no PLK-1 detectable at any of the control centrosomes analyzed at 39±4 seconds post early telophase (n=29 centrosomes) (Figure 3, A and B; Supplemental Figure S3). In contrast, in 35% of the centrosomes from *dyn-1(RNAi)* treated embryos (9 out of 26 centrosomes), PLK-1 was still detectable at the centrosomes at this stage (p=0.0005) (Figure 3, A and B; Supplemental Figure S3).

**FIGURE 3.**
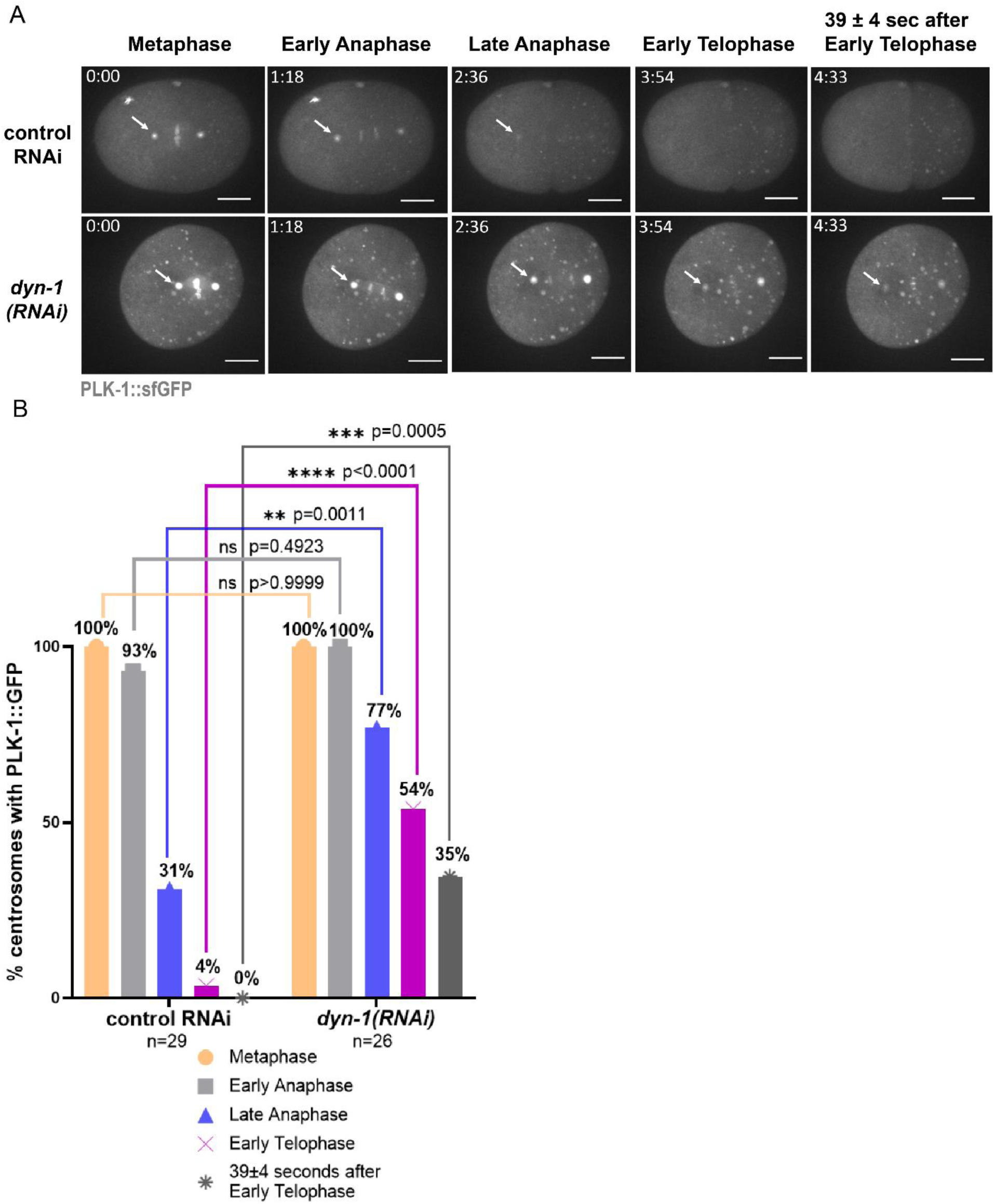
PLK-1 persists abnormally at centrosomes during mitotic progression in *dyn-1(RNAi)* embryos. (A) Time-lapse analysis of different mitotic stages in control RNAi and *dyn-1(RNAi)* embryos. White arrows: anterior centrosome-localized PLK-1::sfGFP. (B) Percentage of centrosomes retaining PLK-1 at each stage. Fisher’s Exact Test; ns= not significant; n = centrosomes analyzed. Experimental replicates: 3 for all assays. All fluorescence images include a 10 µm scale bar.

We also noted a statistically significant increase in centrosome-associated PLK-1::sfGFP upon *dyn-1(RNAi)* (n=26 centrosomes) as compared with controls (n=29 centrosomes) via an unpaired two-tailed t-test at early anaphase (p<0.0001), late anaphase (p=0.0004) as well as in early telophase (p=0.0094) (Supplemental Figure S3). These data indicate that a pool of PLK-1 tends to persist at the centrosomes of DYN-1-depleted embryos during the later stages of mitosis.

### DYN-1-depleted embryos exhibit abnormal midbody formation

Since DYN-1 has a known role in cytokinesis and localizes to the midbody (Thompson *et al.,* 2002), we analyzed midbody formation in DYN-1-depleted embryos. Intercellular bridge (IB) assembly, assessed using mCherry::tubulin fluorescence, served as a proxy for midbody formation (Hu *et al*., 2012; Green *et al.,* 2013; König *et al*., 2017). We noted that all of the control RNAi embryos (n=19 embryos) exhibited a normal tubulin-intense IB during telophase (Figure 4, A (white arrows) and B). In contrast, 50% of the *dyn-1(RNAi)* embryos (12 out of 24 embryos) did not exhibit any detectable IB during telophase (Figure 4, A (bottom panel) and B). Interestingly, 50% of the *dyn-1(RNAi)* embryos exhibited a structurally abnormal IB that either started forming outside the reference time-window for normal IB assembly (normal IB started forming between 195 to 390 seconds post-metaphase) or exhibited defects in IB assembly dynamics (Figure 4, A (middle panel) and B). Importantly, the *dyn-1(ky51)* allele also exhibited mild defects in IB assembly (Figure 4B), demonstrating the specificity of this phenotype. While previous ultrastructural analysis identified a role for DYN-1 in membrane remodeling during the late stages of abscission (König et al., 2017), our study identifies a novel requirement for DYN-1 in regulating midbody organization during the initial stages of this process. Together, these data suggest that DYN-1 function is required for proper midbody assembly.

**FIGURE 4.**
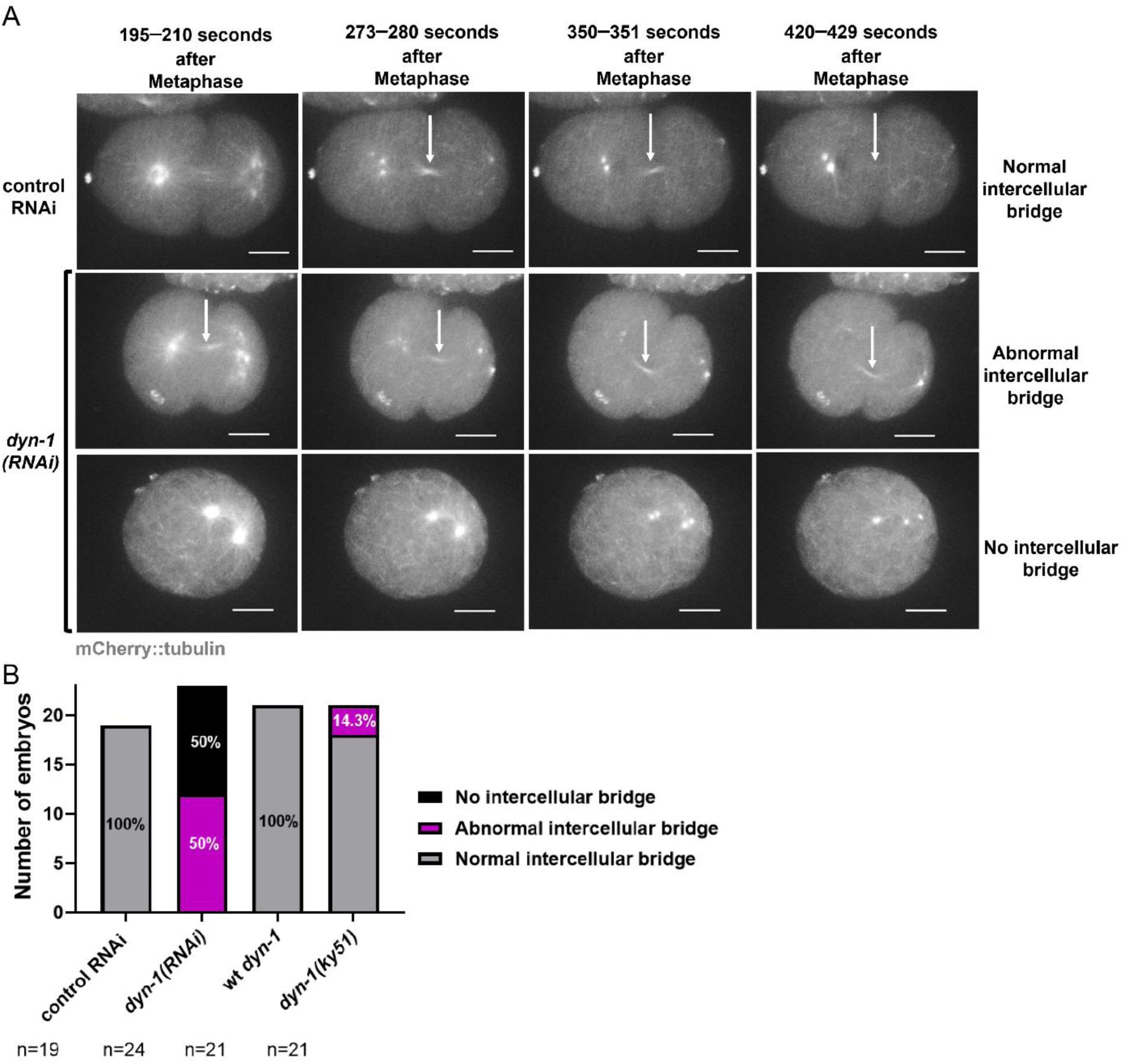
DYN-1 depletion disrupts Intercellular Bridge (IB) assembly. (A) Time-lapse images of mCherry::tubulin showing normal IB formation in control RNAi embryos (top panel) and abnormal (middle panel) or missing (bottom panel) IBs in *dyn-1(RNAi)* embryos. White arrows: IB (B) Classification of IB morphology (normal, abnormal, absent). n = embryos analyzed. Experimental replicates: 3 for all assays. All fluorescence images include a 10 µm scale bar.

### PLK-1 localization to the midbody is defective in DYN-1-depleted embryos

Given that PLK-1 persists at the centrosomes through late mitosis in a subset of DYN-1-depleted embryos (Figure 3), we hypothesized that this retention might correlate with abnormal PLK-1 dynamics at the midbody. To investigate this, we analyzed PLK-1 localization using two complementary approaches: a time-matched analysis and an event-matched analysis. The time-matched analysis (Figure 5, A and B) examined whether PLK-1 localized to midbodies that started forming during the standard developmental window for IB assembly (195–390 seconds post-metaphase). Conversely, the event-matched analysis evaluated PLK-1 recruitment specifically when an IB eventually formed, regardless of timing.

**FIGURE 5.**
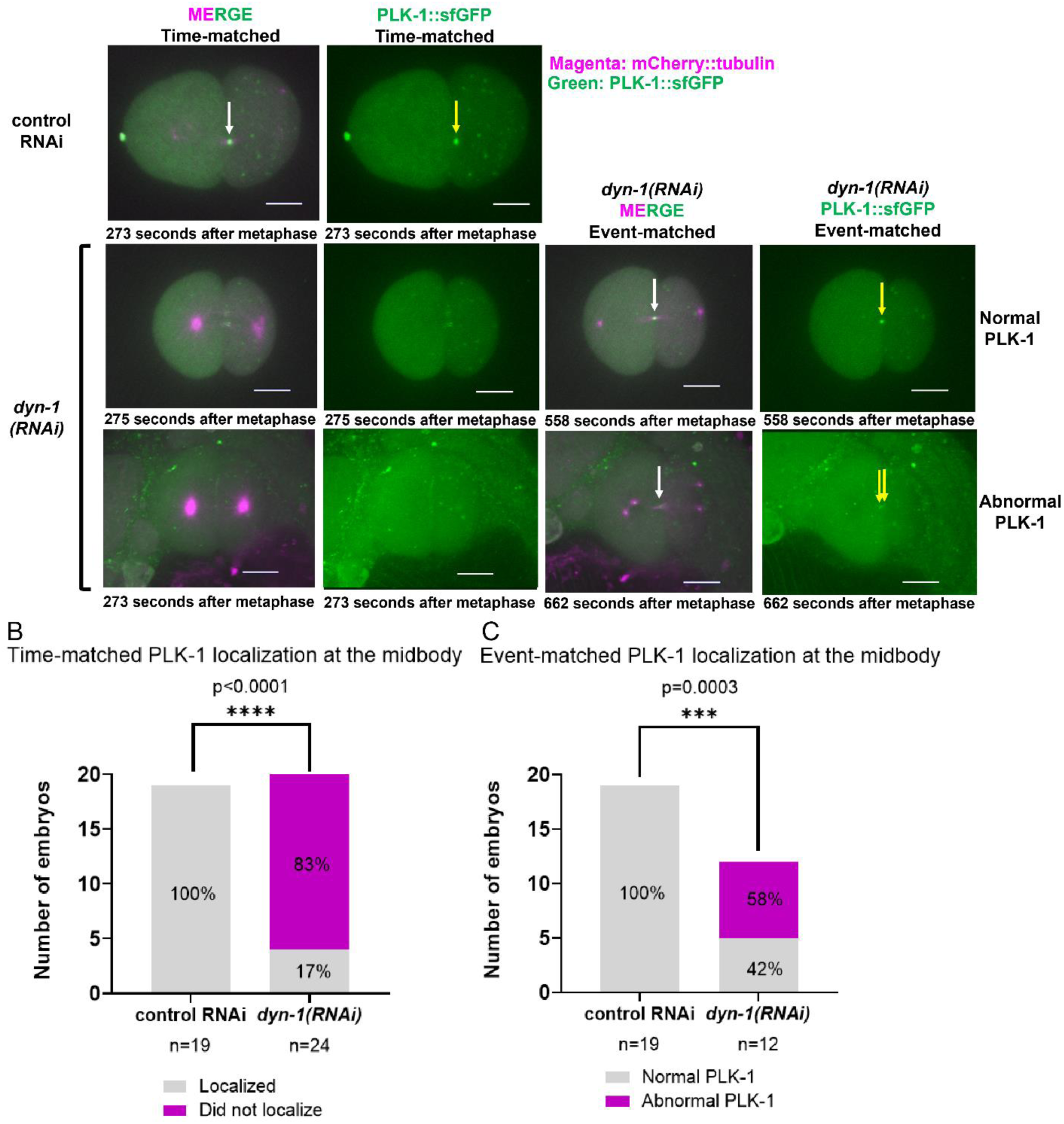
PLK-1 localization at the midbody is defective in *dyn-1(RNAi)* embryos. (A) Time-matched and event-matched PLK-1::sfGFP localization at the midbody in control RNAi and *dyn-1(RNAi)* embryos. White arrows: IB; yellow arrows: PLK-1::sfGFP at IB (B) Time-matched PLK-1 scoring at the midbody. n = embryos; Fisher’s Exact Test. (C) Event-matched PLK-1 scoring at the midbody. n = embryos; Fisher’s Exact Test. Experimental replicates: 3 for all assays. All fluorescence images include a 10 µm scale bar.

Our time-matched analysis revealed that PLK-1 failed to localize to the midbody in 83% of *dyn-1(RNAi)* embryos (n = 24), compared to 100% localization in controls (n = 19; p < 0.0001; Figure 5A). To distinguish whether this reduction was simply due to a failure in midbody formation, we performed an event-matched analysis restricted to *dyn-1(RNAi)* embryos where an IB-like structure was present (n = 12). Importantly, PLK-1 localized to these formed midbodies to varying degrees in all cases: 42% showed relatively normal accumulation, while 58% exhibited fragmented, weak, or near-background signals (Figure 5, A and C; p=0.0003). Together, these data demonstrate that even when a midbody successfully forms following *dyn-1* depletion, PLK-1 recruitment is frequently altered rather than entirely absent. Notably, due to inconsistent background signals resulting from polarity defects in *dyn-1(RNAi)* embryos, we were unable to reliably quantify PLK-1::sfGFP fluorescence intensity at midbodies, as these background fluorescence fluctuations often led to inconsistent measurements.

In summary, our studies have identified a new role for the endocytic protein DYN-1 in regulating metaphase spindle assembly, early midbody assembly and the localization of the mitotic proteins SPD-2 and PLK-1. Specifically, we find that upon DYN-1 depletion, SPD-2 and PLK-1 accumulate at the centrosomes in metaphase and this is associated with aberrant metaphase spindle pole assembly. Furthermore, while PLK-1 is normally removed from the centrosomes during late mitosis, upon DYN-1 depletion, we find that PLK-1 persists at a subset of centrosomes during the later stages of mitosis.

Based upon our data, we propose a model whereby DYN-1 regulates the levels of SPD-2 and PLK-1 at metaphase centrosomes, potentially through a vesicular trafficking mechanism. DYN-1 depletion leads to elevated SPD-2 levels at metaphase centrosomes, and because SPD-2 is a known upstream recruiter of PLK-1 (Boxem *et al*., 2008; Decker *et al*., 2011), this increase in SPD-2 is accompanied by increased PLK-1 accumulation at centrosomes. The elevated levels of these PCM components are associated with aberrant spindle pole assembly. In late mitosis, DYN-1 depletion also disrupts the normal relationship between midbody formation and PLK-1 accumulation at the IB: embryos frequently fail to assemble a midbody, and even in cases where a midbody-like structure does form, PLK-1 enrichment is often fragmented or reduced. While we cannot yet distinguish whether altered PLK-1 midbody localization reflects defective midbody assembly, impaired PLK-1 recruitment, or a combination of both, our findings indicate that DYN-1 is required for coordinating the proper redistribution of PLK-1 between centrosomes and the midbody.

The defects in midbody assembly were more pronounced in *dyn-1(RNAi)* embryos than in *dyn-1(ky51)* mutants. Correspondingly, PLK-1::sfGFP enrichment at metaphase centrosomes was higher following *dyn-1(RNAi)* (∼2.5-fold) compared to *dyn-1(ky51)* (∼1.85-fold). These observations suggest that *dyn-1(ky51)* may be a hypomorphic allele. Alternatively, DYN-1 may regulate these processes through mechanisms other than its role in vesicular trafficking. Future studies should investigate these potential roles, including whether DYN-1 physically interacts with PLK-1 to facilitate its recruitment or turnover. Rescue approaches using hypomorphic *plk-1* alleles may also help clarify the contribution of PLK-1 dysregulation to causality of the observed defects, but such experiments are beyond the scope of the present work. It will also be important to determine whether the DYN-1→SPD-2→PLK-1 pathway is conserved in other organisms.

Since our studies also uncovered the regulation of centrosome-associated SPD-2 levels by DYN-1, an important next step is to determine the specific mechanism of this regulation. Since SPD-2 is an upstream component of the PCM assembly pathway (Kemp *et al*., 2004), future studies will examine whether centrosome levels of other PCM proteins, such as PCMD-1, SAS-7, SPD-5, and AIR-1, are also altered upon DYN-1 depletion. Investigating these broader effects will clarify whether the role of DYN-1 is specific to the SPD-2/PLK-1 axis or if it influences centrosome composition and mitotic spindle assembly more generally.

## MATERIALS AND METHODS

### *C. elegans* strains growth and maintenance

All *C. elegans* strains described in this study were grown and maintained using standard techniques on OP50 seeded Modified Youngren’s Only Bacto-peptone (MYOB) agar plates at 20°C (the strains were shifted to other assay-specific temperatures as needed).

### *C. elegans* strains and genetic crosses

The wild-type N2 strain and the OD2425 strain expressing PLK-1::sfGFP (*plk-1(lt18[plk-1::sfGFP]::loxp) III)* (Martino *et al.,* 2017) were obtained from the *Caenorhabditis Genetics Center* (CGC). The OC908 strain (*bsSi30[pCW9: unc-119(+) pcdk-11.2::sfgfp::his-58::cdk-11.2 3’ UTR] II; bsIs20[pNP99: unc-119(+) tbb-1p::mCherry::tbb-2::tbb-2 3’-UTR]; bsIs2[pCK5.5: Ppie-1::gfp::spd-2]*) and the OC779 strain (*bsSi15[pKO109: spd-2p-spd-2::mCherry::spd-2 3’-utr, unc-119(+)] I; bsSi30[pCW9: unc-119(+) pcdk-11.2::sfgfp::his-58::cdk-11.2 3’ utr] II; unc-119(ed3) III*) were obtained from Dr. Kevin O’Connell’s laboratory at NIDDK, NIH. The mCherry::tubulin–expressing strain IYR028 (*bsIs20[pNP99: unc-119(+) tbb-1p::mCherry::tbb-2::tbb-2 3’-UTR]*) was generated in our laboratory by crossing OC908 hermaphrodites with N2 males and isolating progeny expressing only mCherry::tubulin. Following this, mCherry::tubulin expressing males were generated from the IYR028 strain by heat shock at 30^°^C for 6 hours. These mCherry::tubulin males from the IYR028 strain were then crossed with PLK-1::sfGFP expressing hermaphrodites from the OD2425 strain to generate the IYR038 strain (*bsIs20[pNP99: unc-119(+) tbb-1p::mCherry::tbb-2::tbb-2 3’-UTR]; plk-1(lt18[plk-1::sGFP]::loxP) III)* which was homozygous for both mCherry::tubulin and PLK-1::sfGFP. This strain was validated for homozygosity of the two loci by confocal microscopy. The IYR035 strain (*dyn-1(luv35[ky51]*) *X*) was generated by CRISPR/Cas9 editing as described previously (Paix *et al.,* 2015; Paix *et al.,* 2016; Iyer *et al.,* 2018; Smith *et al*., 2020). This strain was confirmed by both PCR and restriction digestion with Pst1 followed by agarose gel electrophoresis as well as by DNA sequencing. The IYR043 strain (*bsIs20[pNP99: unc-119(+) tbb-1p::mCherry::tbb-2::tbb-2 3′-UTR]; plk-1(lt18[plk-1::sfGFP]::loxP) III; dyn-1(luv35[ky51]) X*) was homozygous for the *dyn-1(ky51)* mutation, mCherry::tubulin and PLK-1::sfGFP. The IYR044 strain (*bsIs20[pNP99: unc-119(+) tbb-1p::mCherry::tbb-2::tbb-2 3′-UTR]; plk-1(lt18[plk-1::sfGFP]::loxP) III; dyn-1(+) X*) was homozygous for mCherry::tubulin, PLK-1::sfGFP and wild-type for *dyn-1*. The IYR043 and IYR044 strains were both generated by crossing IYR035 hermaphrodites with IYR038 males (the males were made by heat shock at 30^°^C for 6 hours). Both these strains were validated by confocal microscopy for homozygosity of the fluorescent markers (mCherry::tubulin and PLK-1::sfGFP). PCR and Pst1 digestion followed by agarose gel electrophoresis as well as by DNA sequencing was used to confirm homozygosity of the *dyn-1(ky51)* mutation (IYR043) and wild-type *dyn-1* (IYR044).

### CRISPR/Cas9 editing

CRISPR/Cas9 genome editing utilizing assembled ribonucleoprotein complexes and a linear DNA repair template (Paix *et al.,* 2015; Paix *et al.,* 2016) was performed to generate the IYR035 strain (*dyn-1(luv35[ky51]*) *X*) according to the protocols detailed in Iyer *et al*. (2018) and Smith *et al*. (2020). The sequences of the editing reagents used to create this strain are as follows:

*dyn-1(ky51)* crRNA: 5’- TGAATAAGCTGCAAAATAAG −3’

*dyn-1(ky51)* repair template (silent mutations introduce a Pst1 restriction site): 5’-AGACTTCTTGCCACGTGGATCAGGAATCGTAACACGTCGTTCATTGATCCTGCAGCTTATTCAAGATCGCAATGAGTACGCCGAGTTCCTAC −3’

*dyn-1(ky51)* screening primers:

*dyn-1(ky51)* forward: 5’- CGTTGATCCCTGTGATCAAT – 3’

*dyn-1(ky51)* reverse: 5’- TTTCAGATGCTCAAATCATCC – 3’

### *RNAi* by feeding

HT115 bacteria expressing the *dyn-1(RNAi)* construct were purchased from Horizon Discovery Ltd. (Lafayette, CO) while HT115 bacteria expressing the *smd-1 RNAi* construct (control) were obtained from Dr. Kevin O’Connell’s laboratory (NIDDK, Bethesda, MD). HT115 *E. coli* transformed with either control (*smd-1(RNAi)*) or *dyn-1(RNAi)* plasmids were grown on MYOB plates supplemented with 100 µg/ml carbenicillin and 2 mM Isopropyl β- d-1-thiogalactopyranoside (IPTG) as described previously (Kamath and Ahringer J 2003; Iyer *et al.,* 2022). Briefly, L4-stage *C. elegans* from the IYR038 strain were transferred onto *RNAi* plates and incubated at 25 °C for 24 to 36 hours. Gravid adults from these plates were then dissected to release their embryos and 1-cell stage embryos were analyzed by confocal live imaging.

### Conditions for wild-type and *dyn-1(ky51)* worms

Starved plates with the IYR043 (*dyn-1(ky51)*) and IYR044 (wild-type *dyn-1*) strains were washed with M9 buffer and used to seed large 10 cm MYOB plates. These plates were placed at 26 °C until L4s appeared. 100 to 125 L4s were picked from each plate and transferred onto new individual 10 cm MYOB plates at 26 °C overnight. Embryos from both strains were imaged the next day using a spinning-disk confocal microscope.

### Sequence of *dyn-1(RNAi)* construct

The commercially purchased *dyn-1(RNAi)* construct (Horizon Discovery Inc.) was subjected to whole plasmid sequencing (Plasmidsaurus). The targeting sequence was found to be as follows: 5’-ATGTCGTGGCAAAACCAGGGAATGCAGGCGTTGATCCCTGTGATCAATCGTGTCCAGGACGCCTTCTCCCAGCTGGGCACATCTGTCAGCTTCGAACTTCCACAGATCGCCGTCGTCGGAGGACAGTCCGCTGGAAAGTCGTCGGTGCTGGAGAATTTTGTCGGAAAAGACTTCTTGCCACGTGGATCAGGAATCGTAACACGTCGTCCACTTATTTTGCAGCTTATTCAAGATCGCAATGAGTACGCCGAGTTCCTACACAAGAAGGGTCATCGCTTTGTGGATTTTGATGCAGTTCGGAAAGAGATTGAGGATGAGACTGATCGTGTCACTGGACAGAATAAGGGAATCAGTCCACATCCAATCAACTTGCGTGTCTTTTCTCCAAATGTTCTAAATCTGACACTCATCGATTTGCCCGGTCTCACAAAAGTGCCCGTCGGAGATCAACCAGCAGATATTGAGCAACAGATCCGTGACATGATTCTCACATTCATCAACCGTGAGACTTGCCTCATTCTTGCCGTCACTCCGGCCAACAGCGATCTCGCCACTTCGGATGCGTTGAAACTTGCGAAGGAAGTCGATCCACAGGGTCTTCGCACGATTGGAGCCCTCACCAAACTTGACTTGATGGACGAGGGAACCGATGCTCGCGAGATCCTCGAGAACAAGCTGTTCACACTTCGTCGTGGCTACGTCGGAGTTGTCAATCGTGGGCAGAAGGATATTGTCGGTCGCAAGGATATTAGAGCTGCTTTGGACGCCGAGAGAAAGTTCTTCATCTCACACCCATCCTACCGACATATGGCTGATCGGTTGGGAACAAGCTACCTTCAACACACTCTTAATCAACAGCTCACCAATCATATCCGTGATACACTGCCAACACTTCGTGATAGTCTTCAAAAGAAGATGTTTGCTATGGAAAAGGATGTGGCCGAGTACAAGAACTACCAGCCAAATGATCCAGGCCGCAAGACCAAGGCTCTTTTGCAAATGGTTACCCAGTTCAATGCTGACATTGAGCGCTCCATTGAAGGTTCCTCTGCAAAGCTGGTTTCAGCCAATGAGCTCAGTGGAGGAGCCCGTATCAATCGGCTTTTCCATGAGCGTTTCCCATTTGAGATTGTTAAAA TGGAAATTGACGAGAAAGAAATGCGCAAAGAAATCCAGTATGCCATCAGAAACATTCACGGTATCCGCGTCGGTCTCTTCACTCCGGATATGGCGTTTGAGGCAATTGCCAAAAAGCAAATCACCCGTCTGAAGGAGCCATCGTTGAAATGCGTTGATCTGGTGGTCAACGAGTTGGCTAATGTGATCAGACAGTGCGCTGACACTATGGCTAGATATCCACGTCTTCGTGACGAGCTGGAAAGAATCGTCGTCTCGCATATGCGTGAACGTGAGCAAATTGCCAAGCAGCAAATTGGGCTCATTGTTGACTACGAACTCGCTTATATGAACACAAACCATGAGGATTTCATTGGATTCAGCAATGCTGAAGCAAAAGCCTCCCAAGGACAATCAGCGAAGAAGAATCTTGGAAACCAGGTGATCAGAAAGGGCTGGCTCTCACTGAGCAACGTATCGTTTGTGCGTGGCTCCAAGGACAATTGGTTTGTGCTCATGTCGGACAGTTTGAGTTGGTACAAAGATGATGAGGAGAAGGAGAAGAAGTACATGCTCCCATTGGATGGTGTCAAGCTGAAGGATATTGAGGGTGGATTTATGTCTCGTAACCACAAGTTTGCTCTGTTCTACCCCGACGGAAAGAACATCTACAAGGATTACAAGCAGCTTGAGTTGGGATGCACCAATTTGGACGAAATTGATGCGTGGAAGGCTTCATTCTTGCGTGCTGGTGTCTATCCAGAAAAGCAGAAGGCACAGGAAGATGAGTCCCAACAAGAGATGGAGGATACCTCGATTGATCCACAACTTGAGAGACAGGTGGAGACAATCCGTAATTTGGTTGATTCCTACATGAGAATCATTACCAAGACAATTAAGGACCTGGTTCCAAAGGCGGTGATGCATTTGATTGTTAACCAGACAGGTGAGTTCATGAAAGATGAACTTTTGGCCCATCTCTACCAATGCGGCGACACTGATGCTCTCATGGAGGAATCTCAAATAGAAGCCCAGAAGCGCGAGGAGATGCTCCGAATGTACCATGCTTGCAAGGAGGCGCTCCGCATTATCTCTGAAGTCAACATGAGCACCCTTGGCGACCAGCCGCCGCCATTGCCGATGTCTGACTACCGCCCACACCCATCTGGACCTTCACCGGTGCCGCGTCCGGCTCCTGCTCCACCAGGCGGACGTCAGGCCCCAATGCCACCACGCGGAGGACCCGGTGCCCCACCACCACCAGG CATGAGACCACCACCAGGTGCGCCAGGAGGCGGCGGTGGCATGTACCCACCGTTGATTCCAAC- 3’

### *dyn-1* RT-PCR

The protocol by Ly *et al.,* 2015 was used to extract RNA from either 50 or 100 control and *dyn-1(RNAi)* treated gravid adult worms. One modification to the protocol involved freezing the worms in the TRIzol reagent (Thermo Fisher Scientific, Catalog #15596018) at –80 °C and storing them until the day of the RNA extraction. RT-PCR was performed using the Qiagen OneStep RT-PCR kit (Qiagen Inc., Catalog # 210210) following the manufacturer’s instructions. The primers used for the RT-PCR were as follows:

*dyn-1* RT-PCR forward: 5’- TGTCGGAAAAGACTTCTTGC – 3’

*dyn-1* RT-PCR reverse: 5’- TGTCAGATTTAGAACATTTGG – 3’

Primers sequences targeting the *act-1* gene (loading control) were obtained from Ly *et al*. (2015).

act1f: 5’- ACGCCAACACTGTTCTTTCC – 3’

act1r: 5’- GATGATCTTGATCTTCATGGTTGA – 3’

Band intensities for all RT-PCR samples were quantified using the open-source software Fiji (Schindelin *et al.,* 2012). Fiji was also used to convert all images to magenta and grayscale.

### *C. elegans* embryo live imaging using a spinning-disk confocal microscope For imaging embryos from the IYR038 and OC779 strains depicted in Figure 1

*C. elegans* embryos obtained from dissected *smd-1(RNAi)* (control) and *dyn-1(RNAi)* worms were imaged using an Olympus Yokagawa X1 spinning-disk confocal microscope as described previously (Bergwell, Smith, *et al*., 2023) with a few alterations. Timelapse imaging was performed using the cellSens software (Olympus America Inc.) by taking 1.5 µm Z-stacks every ∼75 seconds at a 60X magnification with a 1.42 numerical aperture (NA) objective using a Prime 95B CMOS camera. To enable quantitative analysis, all images were captured at the same laser intensity [(50% laser intensity for both 561 nm and 488 nm channels for IYR038) and (60% laser intensity for 561 channel and 35% laser intensity for 488 channel for OC779)] and exposure time (700 msec exposure time for both 561 nm and 488 nm channels for both strains). All brightness and contrast adjustments were applied equally for both the control RNAi and *dyn-1(RNAi)* representative images to enable better image presentation and viewing.

### For imaging embryos from the IYR038, IYR043, and IYR044 strains depicted in Figures 2–5, Supplemental Figure S1, Supplemental Figure S2 and Supplemental Figure S3

A Nikon Ti2-E inverted microscope (Nikon Instruments Inc.) outfitted with a CREST X-Light V3 spinning-disk confocal module, a motorized XY stage with encoder feedback, and a 600-µm Z-piezo drive was used to make time-lapse videos of embryos. Embryos were dissected and mounted on agarose pads, and coverslips were sealed with Vaseline following previously described procedures (Bergwell, Smith, *et al*., 2023). Laser illumination was supplied through a Celesta solid-state light engine (Lumencor Inc.). Time-lapse image acquisition was performed in NIS-Elements (Nikon Instruments Inc.), capturing 1.25-µm optical Z-stacks approximately every 39±4 seconds with a 60× Plan Apo λD oil-immersion objective (NA 1.42; refractive index 1.515) and an ORCA-Fusion BT sCMOS detector (Hamamatsu Inc.). To ensure consistency in quantitative measurements, all datasets were collected under identical exposure conditions: 200 msec exposure and 60% laser power at 561 nm and 400 msec exposure and 35% laser power at 488 nm. Maximum-intensity projections of Z-stacks were generated with NIS-Elements (Nikon Instruments Inc.). For display purposes, brightness and contrast adjustments were applied uniformly across the entire dataset.

### Spindle pole area measurement

To measure spindle pole area, the cellSens software (Olympus America Inc.) was used to obtain maximum intensity Z-projections of the original .vsi files. These Z-projections were saved in a .png format with the fixed scaling (left:80 and right:400) for all images and further analyzed using the Nikon NIS-Elements software (Nikon Instruments Inc.). The calibration on the NIS-Elements software was set to 1 pixel = 0.18333 µm. The thresholding tool in NIS-Elements was used to define and compute the spindle pole area (in µm²) (intense mCherry::tubulin fluorescence demarcated each spindle pole). Only regions with clearly defined spindle poles were included in the analysis. Embryos in which spindle poles could not be reliably distinguished were excluded. Brightness adjustments were equally applied to the images representing spindle pole size in Figure 1A for better presentation and viewing.

### SPD-2 metaphase centrosome intensity measurements

For measuring SPD-2 fluorescence intensity at centrosomes, the raw .vsi files acquired using cellSens software (Olympus America Inc.) were opened in the Fiji software (Schindelin *et al.,* 2012) using the Bio-Formats importer plugin. The global scale was set to 1 pixel = 0.18333 µm. Maximum-intensity Z-projections were generated and used for analyzing SPD-2 fluorescence intensity at metaphase centrosomes. Metaphase was defined as the stage just prior to anaphase onset, when the chromosomes had aligned at the cell center. A region of interest (ROI) measuring 1033.33 pixels² was drawn around each metaphase centrosome and an adjacent area of cytoplasm. The mean SPD-2 fluorescence intensity within each ROI (in arbitrary units, A.U.) was quantified using the measure function in Fiji (Schindelin et al., 2012). Background SPD-2 fluorescence intensity (cytoplasmic ROI) was subtracted from the corresponding centrosomal ROI value to obtain the final background-corrected SPD-2 centrosomal fluorescence intensity. To calculate relative SPD-2 fluorescence levels normalized to control RNAi, all background-corrected centrosomal fluorescence intensity values from both control RNAi and *dyn-1(RNAi)* were divided by the mean background-corrected SPD-2 fluorescence intensity measured in control RNAi embryos. Brightness, contrast, and sharpness adjustments were uniformly applied to all images shown in Figure 1C to improve visualization of SPD-2 centrosome localization. Embryos where the Z-projections did not capture the entire centrosome or where SPD-2 at the centrosome could not be clearly identified due to surrounding debris were excluded from our analysis.

### PLK-1 metaphase centrosome intensity measurements

Metaphase was defined as the stage when the chromosomes had aligned at the center of the metaphase plate. To quantify PLK-1::sfGFP fluorescence at the centrosomes, maximum intensity Z-projections of metaphase embryos were generated and analyzed using Nikon NIS-Elements software (Nikon Instruments Inc.). A region of interest (ROI) measuring 11.62 µm² was drawn around each centrosome and within the cytoplasm of the same embryo. Cytoplasmic PLK-1::sfGFP intensity was then subtracted from centrosomal PLK-1::sfGFP intensity to calculate the background-corrected PLK-1::sfGFP centrosome fluorescence intensity, reported in arbitrary units (A.U.). All background-corrected PLK-1::sfGFP centrosome intensities were divided by the mean background-corrected PLK-1::sfGFP centrosome intensity of control embryos to calculate the relative PLK-1 levels at centrosomes. Embryos with inaccurate PLK-1::sfGFP fluorescence due to debris on top of the embryo affecting maximum intensity Z-projections or instances where the Z-projections of the embryo did not cover the entire centrosome were excluded from our analysis.

### Analysis of PLK-1 persistence at centrosomes

To analyze PLK-1::sfGFP persistence at centrosomes at different cell cycle stages, maximum intensity Z-projections of each timelapse sequence were quantified for centrosome-localized PLK-1::sfGFP using the Nikon NIS-Elements software (Nikon Instruments Inc.). To quantify centrosomal PLK-1::sfGFP levels across different cell-cycle stages, measurements were restricted to the anterior centrosome. In the anterior cytoplasm, PLK-1::sfGFP background fluorescence was relatively uniform, enabling consistent background subtraction and accurate intensity quantification. By contrast, the posterior cytoplasm frequently exhibited irregular, punctate PLK-1::sfGFP foci of variable sizes and intensities, which introduced inconsistency in centrosome fluorescence measurements, especially at later cell cycle stages. For this reason, posterior centrosomes were excluded from our quantitative analysis. The microtubule asters were used to identify the position of the centrosomes at each cell cycle stage. A ROI of 11.62 µm² was placed at the center of the asters as well as in its immediate vicinity (background). The PLK-1::sfGFP fluorescence intensity at each anterior centrosome at every cell cycle stage was determined by subtracting the background fluorescence intensity from the centrosomal fluorescence intensity at that stage. Centrosomes with background-corrected PLK-1::sfGFP fluorescence intensities greater than 5 arbitrary units (A.U.) were classified as “PLK-1 present” whereas centrosomes with intensity values below this threshold were classified as “PLK-1 absent”. The cell cycle stages were classified as follows: metaphase: a diamond-shaped spindle formed and chromosomes aligned at the center, early anaphase: ∼39±4 seconds to 78±8 seconds after metaphase when the chromosomes started separating, late anaphase: ∼78±8 seconds after early anaphase, early telophase: ∼78±8 seconds after late anaphase, and 39±4 seconds after early telophase: ∼39±4 seconds after early telophase. In the occasional cases where PCM disassembly prevented identification of the aster center, the centrosome position from the preceding frame was used as a reference. Brightness adjustments were applied equally to the representative images in Figure 3A for better presentation and viewing. Only embryos where the Z-projections captured the entire centrosome were used for our analysis.

### Quantification of midbody abnormalities

Maximum-intensity Z-projections generated using Nikon NIS-Elements (Nikon Instruments Inc.) were used to analyze midbody defects. Embryos were scored in a randomized and blinded manner: image files were renamed with randomized identifiers and scored by an independent individual. The scorer evaluated IB behavior without knowledge of genotype or treatment until scoring was complete. An intercellular bridge (IB) was defined as present when a continuous, tubulin-intense connection between the two daughter cells could be clearly visualized. IB assembly was used as an indicator of midbody formation (Hu *et al*., 2012; Green *et al*., 2013; König *et al*., 2017).

A normal IB was defined as one that started forming approximately 195–390 seconds after metaphase onset, remained stable for at least 105 seconds following its formation, and exhibited the stereotypical IB assembly dynamics. Stereotypical IB dynamics consisted of a sequence in which the spindle midzone coalesced into a single focused, microtubule-rich bridge-like IB structure. Following coalescence, the IB typically increased in tubulin intensity, then gradually decreased in intensity as it regressed. Once coalesced, normal IBs did not separate or fragment and maintained a continuous tubulin-dense morphology until midbody resolution.

An abnormal IB was defined as one that failed to exhibit this stereotypical progression. Abnormal IBs included bridges that did not start forming within the 195–390 second window, did not coalesce into a single focused structure, showed fluctuating tubulin intensity, or came apart after coalescence. An embryo was scored as lacking an IB when no stable IB could be detected at any point during the remainder of the cell cycle and through the onset of centrosome separation in the next cell cycle.

Embryos where the entire IB was not captured in the maximum intensity Z-projections were excluded from our analysis. For better visualization and presentation, brightness and contrast setting were applied uniformly and equally to all the representative images in Figure 4A.

### Quantification of PLK-1 localization at the midbody

PLK-1 localization at the midbody was quantified using a blinded, randomized analysis using maximum intensity Z-projections that were generated with the Nikon NIS-Elements software (Nikon Instruments Inc.). All image files were renamed with random alphanumeric identifiers by an independent individual who was not involved in the scoring. The scorer analyzed the randomized dataset without knowledge of genotype, treatment condition, or imaging sequence. After all scoring was completed and recorded, the mapping key was revealed to assign quantified values back to experimental groups.

An IB/midbody was defined as present when a continuous, tubulin-intense connection between the two daughter cells could be clearly visualized in Z-projections. An IB was considered to have formed if it remained cohesive for at least ∼105 seconds after its appearance. Metaphase was defined as the timepoint immediately after nuclear envelope breakdown when the chromosomes are aligned centrally at the metaphase plate. For event-matched analysis, a normal PLK-1 localization pattern was defined as a continuous, dot-like or band-like PLK-1::sfGFP signal above the local background at the IB center. Abnormal localization was defined as discontinuous, fragmented, or off-center PLK-1::sfGFP fluorescence, or a reduced signal that was either at or below the local background. For the time-matched analysis, cases where PLK-1::sfGFP localized to the IB were scored as “localized” and those where it failed to localize to the IB either because IB formation was delayed or because an IB did not form were scored as “did not localize”. For all analyses, PLK-1 localization was scored at the frame in which the IB was fully formed, and just before the tubulin intensity at the IB began to decrease, ensuring that measurements were taken while the midbody remained structurally relatively intact.

For time-matched analysis, PLK-1::sfGFP fluorescence was assessed relative to the reference window during which stable IB formation normally begins in control embryos (195–390 seconds after metaphase onset).

For event-matched analysis, PLK-1::sfGFP fluorescence was assessed whenever a stable IB formed, even if the IB formed outside of the normal developmental window for bridge assembly.

Z-projections in which either the IB or midbody-associated PLK-1 were not fully captured across the Z-stack were excluded from analysis to ensure accurate evaluation of PLK-1 fluorescence at the IB/midbody. Brightness adjustments were applied equally to all merged and all PLK-1::sfGFP representative images of PLK-1 midbody localization in Figure 5A.

### Embryo size measurements

Nikon-NIS Elements (Nikon Instruments Inc.) was used to generate maximum intensity Z-projections of the raw data and to measure the sizes of metaphase-stage 1-cell *C. elegans* embryos. To measure embryo size, the Bézier tool in NIS-Elements was used to manually trace the outline of each embryo, using cytoplasmic PLK-1::sfGFP fluorescence to define the cytoplasmic embryo boundary. The enclosed area was calculated by the software and reported in µm². Embryos whose boundaries could not be clearly defined by cytoplasmic PLK-1::sfGFP fluorescence due to embryo orientation were excluded from the analysis.

### Cytoplasmic PLK-1 fluorescence measurements

Maximum intensity Z-projections generated by Nikon NIS-Elements (Nikon Instruments Inc.) were used to measure cytoplasmic PLK-1::sfGFP in the anterior and posterior cytoplasm of wild-type and *dyn-1(ky51)* embryos. For these measurements, the Z-projections were opened in NIS-Elements and a large ROI of 128.97 µm^2^ was placed within the anterior and posterior cytoplasm of metaphase-stage embryos away from the spindle apparatus. The mean PLK-1::sfGFP fluorescence intensities within these ROIs were determined using the NIS-Elements software and reported in arbitrary units (A.U.). Measurements in which the ROI could not fit within the embryo boundaries due to smaller embryo size were excluded from our analysis.

### Statistical analysis

Graphpad prism 10.5 and 10.6.1 (GraphPad Software, Inc., San Diego, CA) were used to perform statistical analyses of all the data. For all the data, error bars represent the standard deviation (s.d.) above and below the mean and the middle lines represent the mean. *p*-values were rounded to two decimal places and considered significant if *p*<0.05. A two-tailed unpaired t-test was used to analyze: spindle pole area (Figure 1B), SPD-2 fluorescence intensity (Figure 1D), PLK-1::sfGFP centrosome fluorescence intensities at different cell cycle stages (Supplemental Figure S3), *dyn-1* knockdown by RT-PCR (Supplemental Figure S1B), embryo size (cytoplasmic area) (Supplemental Figure S1D), and cytoplasmic PLK-sfGFP intensity (Supplemental Figure S2). A Fisher’s exact test was used to analyze the PLK-1::sfGFP persistence at the centrosomes (Figure 3B), and defects in PLK-1::sfGFP localization to the midbody (Figure 5, B and C).

## ACKNOWLEDGMENTS

This work was supported by NIH/NIGMS grants (5P20GM103636-09 and 7R15GM152965-02) and start-up funding from North Carolina A&T State University to J.I. Additional support was provided by NIH/NIGMS grant 1R35GM153737 and NCI grant 1R01CA273095 to R.N. We would like to acknowledge Dr. Hailey Edigo-Betancourt, Jasmyn Grant, Journey Reddick-Umoja, Erin Haastrup, Peter Nunan, and Mariah Moreland for technical assistance. OC908 and OC779 strains were provided by Dr. Kevin O’Connell (NIDDK, NIH). Some strains used in this study were provided by the CGC, which is funded by NIH Office of Research Infrastructure Programs (P40 OD010440). The authors declare no competing interests.

## SUPPLEMENTAL FIGURES

FIGURE S1. *dyn-1(RNAi)* reduces *dyn-1* mRNA levels and decreases embryo size. (A) RT-PCR validation of *dyn-1* depletion. (B) Relative *dyn-1* mRNA quantification. Error bars = s.d. (C) Comparison of average embryo sizes in control RNAi and *dyn-1(RNAi)*. (D) Cytoplasmic area quantification using PLK-1::sfGFP fluorescence as a marker for the cytoplasmic embryo border. n = embryos. Error bars = s.d.; middle bar = mean. (B, D): unpaired two-tailed t-test; Experimental replicates: 3 for all assays. All fluorescence images include a 10 µm scale bar.

FIGURE S2. Cytoplasmic PLK-1::sfGFP levels are modestly elevated in *dyn-1(ky51)* embryos as compared to wild-type. (A) Quantification of anterior and posterior cytoplasmic PLK-1::sfGFP in wild-type (wt) *dyn-1* and *dyn-1(ky51)* embryos. A.U.= Arbitrary Units; unpaired two-tailed t-test; n = measurements. Error bars = s.d.; middle bar = mean. Experimental replicates: 3.

FIGURE S3. Comparison of centrosomal PLK-1 intensity across cell cycle stages in control RNAi and *dyn-1(RNAi)* embryos. (A) Centrosomal PLK-1::sfGFP intensity at metaphase, early and late anaphase, early telophase, and 39±4 seconds post early telophase. A.U.= Arbitrary Units; unpaired two-tailed t-test; n = 29 centrosomes (control RNAi); n = 26 centrosomes (*dyn-1(RNAi))*. Error bars = s.d.; middle bar = mean. Experimental replicates: 3.

## REFERENCES

Archambault V, Glover DM (2009). Polo-like kinases: conservation and divergence in their functions and regulation. Nat Rev Mol Cell Biol 10, 265–275.

Arnaud L, Pines J, Nigg EA (1998). GFP tagging reveals human Polo-like kinase 1 at the kinetochore/centromere region of mitotic chromosomes. Chromosoma 107, 424–429.

Barr FA, Silljé HHW, Nigg EA (2004). Polo-like kinases and the orchestration of cell division. Nat Rev Mol Cell Biol 5, 429–441.

Bergwell M, Smith A, Smith E, Dierlam C, Duran R, Haastrup E, Napier-Jameson R, Seidel R, Potter W, Norris A, et al. (2023). A primary microcephaly-associated *sas-6* mutation perturbs centrosome duplication, dendrite morphogenesis, and ciliogenesis in *Caenorhabditis elegans*. Genetics 224, iyad105.

Boxem M, Maliga Z, Klitgord N, Li N, Lemmens I, Mana M, De Lichtervelde L, Mul JD, Van De Peut D, Devos M, et al. (2008). A protein domain-based interactome network for *C. elegans* early embryogenesis. Cell 134, 534–545.

Budirahardja Y, Gönczy P (2008). PLK-1 asymmetry contributes to asynchronous cell division of *Caenorhabditis elegans* embryos. Development 135, 1303–1313.

Chase D, Serafinas C, Ashcroft N, Kosinski M, Longo D, Ferris DK, Golden A (2000). The polo-like kinase PLK-1 is required for nuclear envelope breakdown and the completion of meiosis in *Caenorhabditis elegans*. Genesis 26, 26–41.

Chen YZ, Zimyanin V, Redemann S (2023). Loss of the mitochondrial protein SPD-3 elevates PLK-1 levels and dysregulates mitotic events. Life Sci Alliance 6, e202302117.

Clark SG, Shurland DL, Meyerowitz EM, Bargmann CI, Van der Bliek AM (1997). A dynamin GTPase mutation causes a rapid and reversible temperature-inducible locomotion defect in *Caenorhabditis elegans*. Proc Natl Acad Sci USA 94, 10438–10443.

Decker M, Jaensch S, Pozniakovsky A, Zinke A, O’Connell KF, Zachariae W, Myers E, and Hyman AA (2011). Limiting amounts of centrosome material set centrosome size in *C. elegans* embryos. Curr Biol 21, 1259–1267.

Golsteyn RM, Mundt KE, Fry AM, Nigg EA (1995). Cell cycle regulation of the activity and subcellular localization of Plk1, a human protein kinase implicated in mitotic spindle function. J Cell Biol 129, 1617–1628.

Gomez-Ferreria MA, Rath U, Buster DW, Chanda SK, Caldwell JS, Rines DR, Sharp DJ (2007). Human Cep192 is required for mitotic centrosome and spindle assembly. Curr Biol 17, 1960–1966.

Grant B, Hirsh D (1999). Receptor-mediated endocytosis in the *Caenorhabditis elegans* oocyte. Mol Biol Cell 10, 4311–4326.

Green, R.A., J.R. Mayers, S. Wang, L. Lewellyn, A. Desai, A. Audhya, and K. Oegema (2013). The midbody ring scaffolds the abscission machinery in the absence of midbody microtubules. J Cell Biol. 203, 505–520.

Hinshaw JE (2000). Dynamin and its role in membrane fission. Annu Rev Cell Dev Biol 16, 483–519.

Hu CK, Coughlin M, Mitchison TJ (2012). Midbody assembly and its regulation during cytokinesis. Mol Biol Cell 23, 1024–1034.

Iyer J, Gentry LK, Bergwell M, Smith A, Guagliardo S, Kropp PA, Sankaralingam P, Liu Y, Spooner E, Bowerman B, et al. (2022). The chromatin remodeling protein CHD-1 and the EFL-1/DPL-1 transcription factor cooperatively downregulate CDK-2 to control SAS-6 levels and centriole number. PLoS Genet 18, e1009799.

Iyer J, DeVaul N, Hansen T, Nebenfuehr B (2019). Using microinjection to generate genetically modified *Caenorhabditis elegans* by CRISPR/Cas9 editing. In: Microinjection: Methods and Protocols, ed. C Liu and Y Du, New York, NY: Humana Press, 431–457.

Kamath RS, Ahringer J (2003). Genome-wide RNAi screening in *Caenorhabditis elegans*. Methods 30, 313–321.

Kemp CA, Kopish KR, Zipperlen P, Ahringer J, O’Connell KF (2004). Centrosome maturation and duplication in *C. elegans* require the coiled-coil protein SPD-2. Dev Cell 6, 511–523.

Kishi K, Van Vugt MA, Okamoto KI, Hayashi Y, Yaffe MB (2009). Functional dynamics of Polo-like kinase 1 at the centrosome. Mol Cell Biol 29, 3134–3150.

König J, Frankel EB, Audhya A, Müller-Reichert T (2017). Membrane remodeling during embryonic abscission in *Caenorhabditis elegans*. J Cell Biol 216, 1277–1286.

Lee KS, Yuan YL, Kuriyama R, Erikson RL (1995). Plk is an M-phase-specific protein kinase and interacts with a kinesin-like protein, CHO1/MKLP1. Mol Cell Biol 15, 7143–7151.

Levine MS, Holland AJ (2018). Impact of mitotic errors on cell proliferation and tumorigenesis. Genes Dev 32, 620–638.

Ly K, Reid SJ, Snell RG (2015). Rapid RNA analysis of individual *Caenorhabditis elegans*. MethodsX 2, 59–63.

Martino L, Morchoisne Bolhy S, Cheerambathur DK, Van Hove L, Dumont J, Joly N, Desai A, Doye V, Pintard L (2017). Channel nucleoporins recruit PLK-1 to nuclear pore complexes to direct nuclear envelope breakdown in *C. elegans*. Dev Cell 43, 157–171.

O’Connell KF, Maxwell KN, White JG (2000). The *spd-2* gene is required for axis polarization in the *C. elegans* zygote. Dev Biol 222, 55–70.

Paix A, Folkmann A, Rasoloson D, Seydoux G (2015). High efficiency, homology-directed genome editing in Caenorhabditis elegans using CRISPR-Cas9 ribonucleoprotein complexes. Genetics 201, 47–54.

Paix A, Schmidt H, Seydoux G (2016). Cas9-assisted recombineering in *C. elegans*: genome editing using in vivo assembly of linear DNAs. Nucleic Acids Res 44, e128.

Paschal BM, Shpetner HS, Vallee RB (1987). MAP 1C is a microtubule-activated ATPase which translocates microtubules in vitro and has dynein-like properties. J Cell Biol 105, 1273–1282.

Pelletier L, Özlü N, Hannak E, Cowan C, Habermann B, Ruer M, Müller-Reichert T, Hyman AA (2004). The *Caenorhabditis elegans* centrosomal protein SPD-2 is required for both pericentriolar material recruitment and centriole duplication. Curr Biol 14, 863–873.

Petronczki M, Lénárt P, Peters JM (2008). Polo on the rise—from mitotic entry to cytokinesis with Plk1. Dev Cell 14, 646–659.

Praefcke GJA, McMahon HT (2004). The dynamin superfamily: universal membrane tubulation and fission molecules?. Nat Rev Mol Cell Biol 5, 133–147.

Schindelin J, Arganda-Carreras I, Frise E, Kaynig V, Longair M, Pietzsch T, Preibisch S, Rueden C, Saalfeld S, Schmid B, Tinevez J-Y, et al. (2012). Fiji: an open-source platform for biological-image analysis. Nat Methods 9, 676–682.

Schmucker S, Sumara I (2014). Molecular dynamics of PLK1 during mitosis. Mol Cell Oncol 1, e954507.

Shpetner HS, Vallee RB (1989). Identification of dynamin, a novel mechanochemical enzyme that mediates interactions between microtubules. Cell 59, 421–432.

Smith A, Bergwell M, Smith E, Mathew D, Iyer J (2020). CRISPR/Cas9 editing of the *C. elegans rbm-3.2* gene using the *dpy-10* Co-CRISPR screening marker and assembled ribonucleoprotein complexes. J Vis Exp 166, e62001.

Sugioka K, Hamill DR, Lowry JB, McNeely ME, Enrick M, Richter AC, Kiebler LE, Priess JR, Bowerman B (2017). Centriolar SAS-7 acts upstream of SPD-2 to regulate centriole assembly and pericentriolar material formation. eLife 6, e20353.

Thompson HM, Skop AR, Euteneuer U, Meyer BJ, McNiven MA (2002). The large GTPase dynamin associates with the spindle midzone and is required for cytokinesis. Curr Biol 12, 2111–2117.

Zhu F, Lawo S, Bird A, Pinchev D, Ralph A, Richter C, Müller-Reichert T, Kittler R, Hyman AA, Pelletier L (2008). The mammalian SPD-2 ortholog Cep192 regulates centrosome biogenesis. Curr Biol 18, 136–141.

Zitouni S, Nabais C, Jana SC, Guerrero A, Bettencourt-Dias M (2014). Polo-like kinases: structural variations lead to multiple functions. Nat Rev Mol Cell Biol 15, 433–452.

