## Supplementary figures and images for "DYN-1 regulates SPD-2 and PLK-1 localization and mitotic spindle pole organization"

### Supplemental Figures S1-S3

**Supplemental Figure S1**

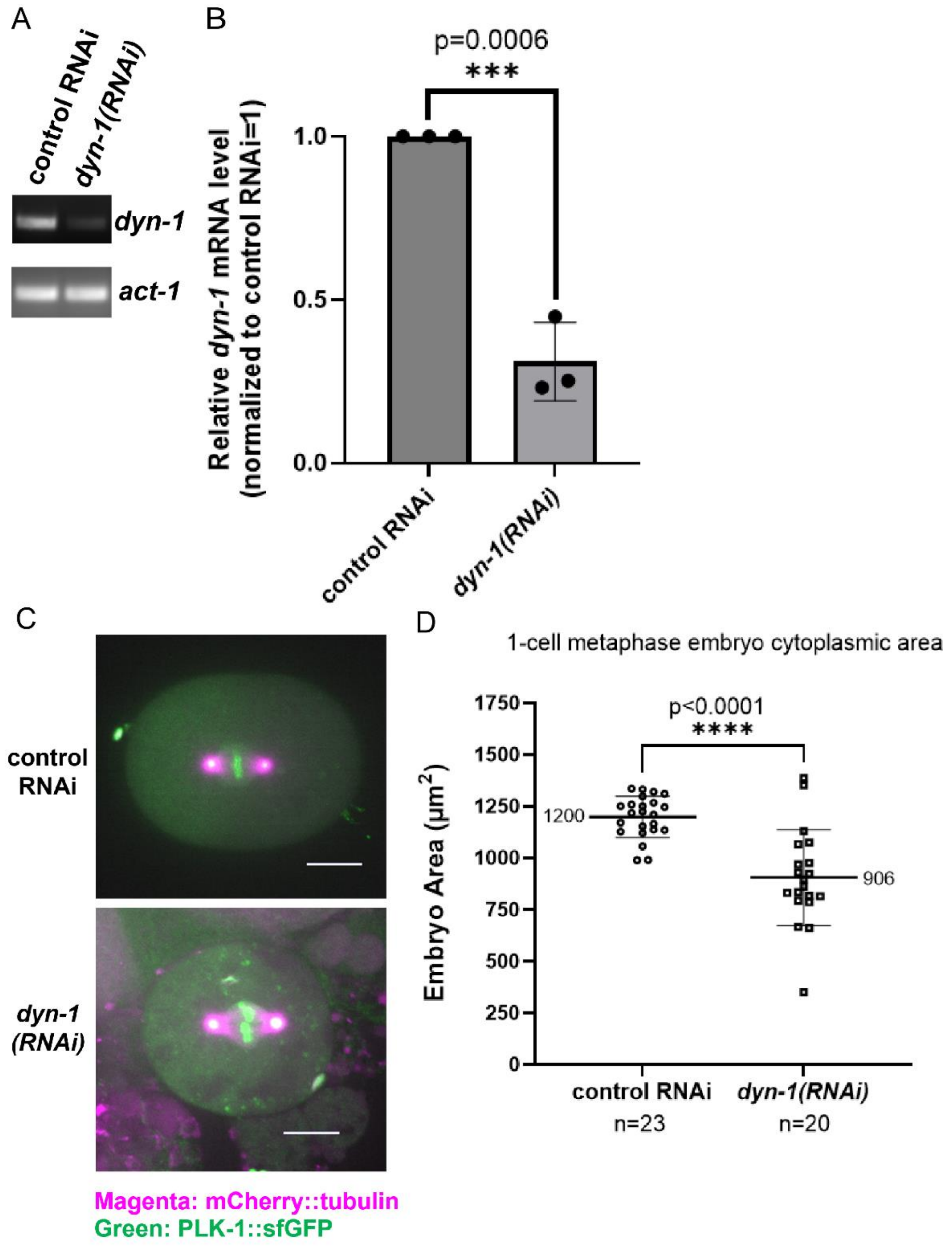

#### Supplemental Figure S2

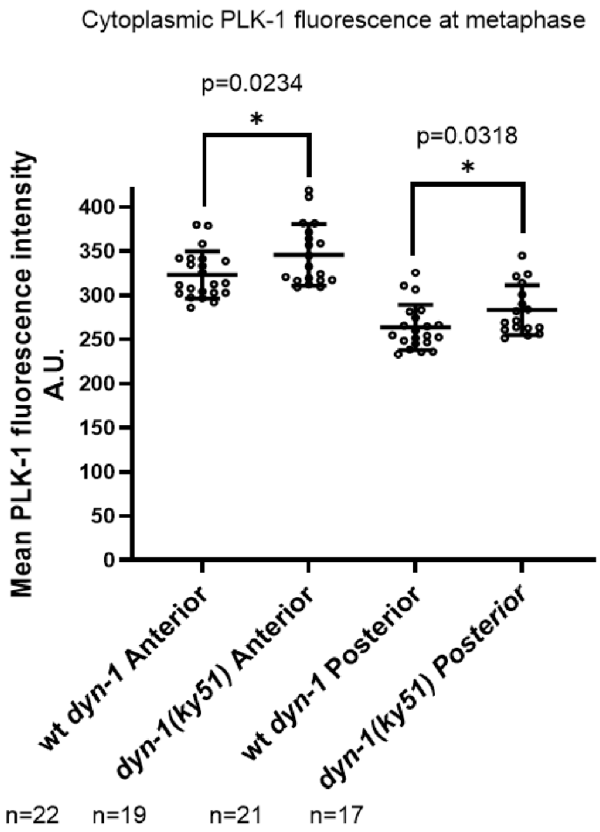

**Supplemental Figure S3**

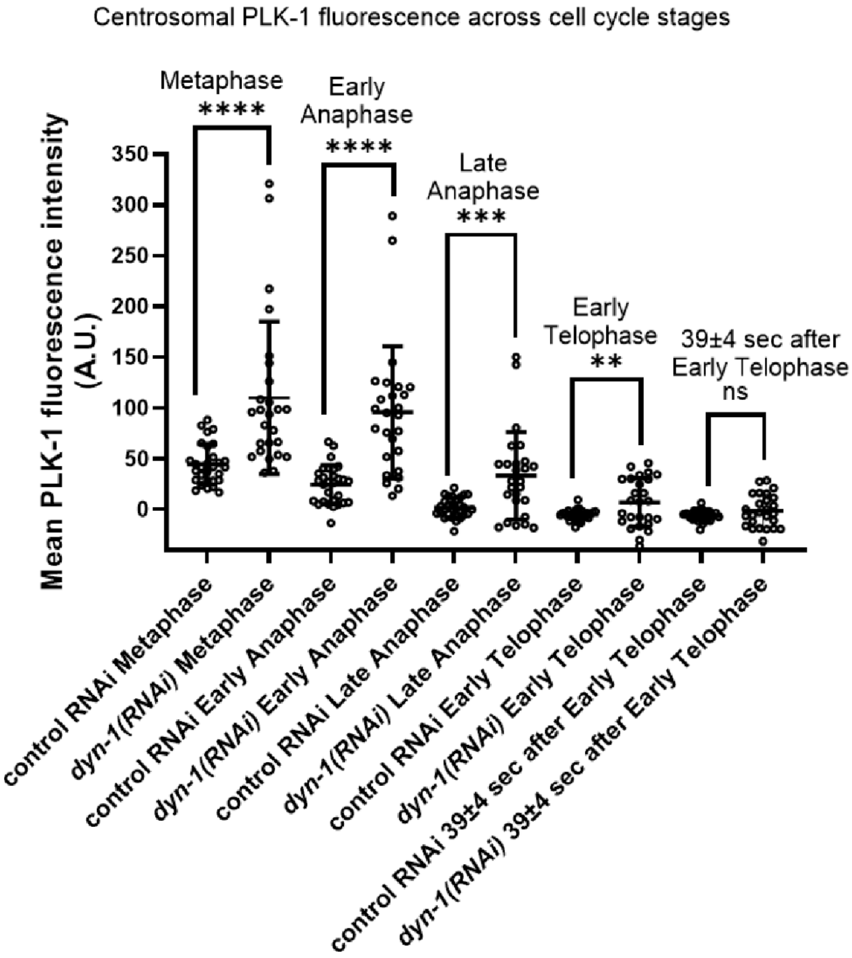
